# Considerations in using recurrent neural networks to probe neural dynamics

**DOI:** 10.1101/364489

**Authors:** Jonathan C Kao

## Abstract

Recurrent neural networks (RNNs) are increasingly being used to model complex cognitive and motor tasks performed by behaving animals. Here, RNNs are trained to reproduce animal behavior while also recapitulating key statistics of empirically recorded neural activity. In this manner, the RNN can be viewed as an *in silico* circuit whose computational elements share similar motifs with the cortical area it is modeling. Further, as the RNN’s governing equations and parameters are fully known, they can be analyzed to propose hypotheses for how neural populations compute. In this context, we present important considerations when using RNNs to model motor behavior in a delayed reach task. First, by varying the network’s nonlinear activation and rate regularization, we show that RNNs reproducing single neuron firing rate motifs may not adequately capture important population motifs. Second, by visualizing the RNN’s dynamics in low-dimensional projections, we demonstrate that even when RNNs recapitulate key neurophysiological features on both the single neuron and population levels, it can do so through distinctly different dynamical mechanisms. To militate between these mechanisms, we show that an RNN consistent with a previously proposed dynamical mechanism is more robust to noise. Finally, we show that these dynamics are sufficient for the RNN to generalize to a target switch task it was not trained on. Together, these results emphasize important considerations when using RNN models to probe neural dynamics.

## Introduction

Recurrent neural networks (RNNs) have been employed to model computation in neurophysiological tasks (Mante et al., 2013; Hennequin et al., 2014; Sussillo et al., 2015; Michaels et al., 2016; Song et al., 2016; Chaisangmongkon et al., 2017; Miconi, 2017; Song et al., 2017). In these studies, the RNN is trained to perform tasks and reproduce empirically observed behavior. Examples include an animal’s kinematics or electromyography during a motor task or its psychometric curve during a decision making task. Further, the RNN can be trained so that its artificial neurons recapitulate key statistics of neurons recorded from experiments, both on the single unit and population level. Training techniques to achieve this include regularizing the network to avoid complex patterns (Sussillo et al., 2015; Michaels et al., 2016), introducing architectural constraints such as Dale’s law (Song et al., 2016), and utilizing biologically plausible learning rules including those based on reinforcement learning (Miconi, 2017; Song et al., 2017). RNNs that recapitulate the behavior and key statistics of the neural population have then been analyzed to propose mechanisms for how recurrent computation occurs in cortical circuits (Sussillo & Barak, 2013; Mante et al., 2013; Chaisangmongkon et al., 2017). The RNN may also generate hypotheses that can be tested in future neurophysiological experiments (Chandrasekaran, 2017).

The existence of a diversity of training approaches that meaningfully change artificial neuron motifs raises several questions. For example, does the particular training approach matter? Said differently, can a variety of RNNs, each trained in a different way but nevertheless all resembling empirical neural activity, employ different dynamical mechanisms? If so, what are the key considerations in using RNNs as *in silico* models of cortical circuits? We address these questions by changing various design variables for RNNs and assessing how these changes affect the RNN’s motifs and dynamics. In particular, we vary (1) the nonlinear activation of the RNN, (2) rate regularization during training, and (3) task input configuration. We perform these comparisons for a common motor neuroscience task: the delayed reach task. We chose this task because prior work in motor systems neuroscience has proposed a concrete dynamical mechanism employed by the motor cortex to perform this task (Ames et al., 2014; Churchland et al., 2006; Afshar et al., 2011; Kaufman et al., 2014). Therefore, we are able to make comparisons to neurophysiological results at the level of the RNN’s single units, population, and dynamics.

We consider how design choices affect the RNN’s ability to recapitulate key behavioral and neural features from experiments. From this, we find that is important to recapitulate both single unit and population motifs. That is, it is possible to find artificial neurons that resemble single unit peri-stimulus time histograms (PSTHs) but that do not capture key population features in the neurophysiological data. Further, we show that distinct RNNs can resemble neurophysiological data while using fundamentally different dynamical mechanisms. We illustrate this idea in Fig 1b-d, where for the same neural population activity evolving in two dimensions, different dynamics (denoted by the flow fields) may give rise to the same population activity. By visualizing the RNN’s dynamical equations, we demonstrate that RNN input design can substantially modify the network’s dynamical mechanisms. Finally, we explore consequences of computation using these distinct dynamical mechanisms, including robustness to noise and generalization to new tasks.

**Figure 1:**
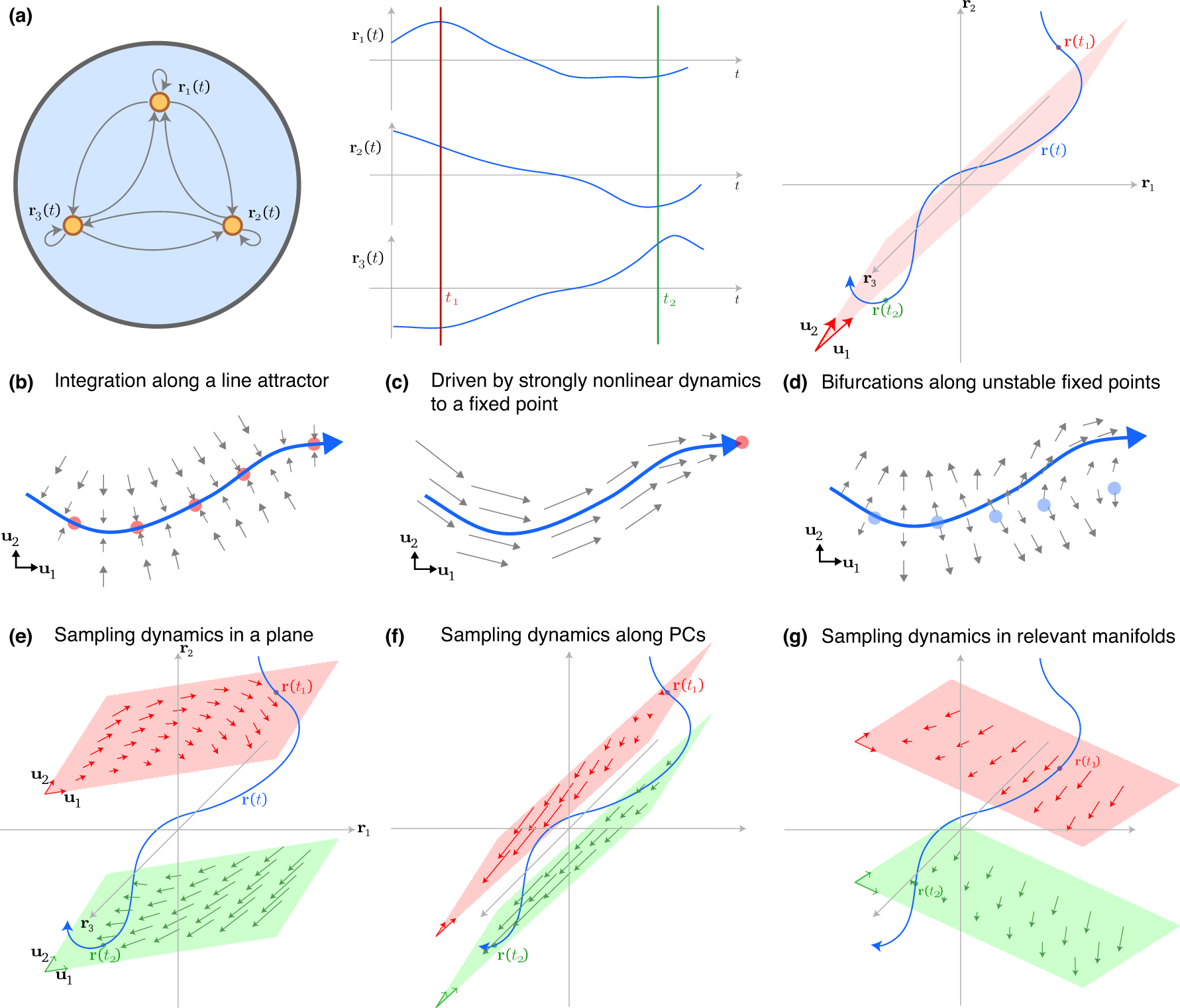
Illustration of sampling RNN dynamics. **(a)** Toy example of an RNN with three example units. The units’ firing rates through time, **r**(*t*), is plotted as a trajectory (blue) in 3 dimensions, but it largely evolves in a 2 dimensional plane indicated in red, defined by the vectors **u**_1_ and **u**_2_. **(b)** The inputs may drive the trajectory slowly along a line attractor. Red dots denote stable fixed points. **(c)** The dynamics may be strong and cause the trajectory to be strongly driven to a stable fixed point. **(d)** The trajectory may be driven along regions of bifurcation, with slow unstable attractors denoted by blue dots. **(e)** For a given basis, defined by **u**_1_ and **u**_2_, it is possible to project the RNN dynamics into a given plane. Here, we show a sampling rule where the values in orthogonal dimensions are set to the trajectory values. An obstacle is that the trajectories sampled at two different times, *t*_1_ and *t*_2_, may have very different dynamics, indicated by the flow field arrows in the red and green planes. **(f)** If the dynamics are relatively smooth, one strategy to address this obstacle is to ensure the sampling planes, shown in red and green, are close to each other. This is achieved by sampling the principal components. **(g)** Another approach is to sample dynamics in “dynamics relevant” manifolds, where the views of the dynamics may not change as drastically depending on the sampling.

## Materials and Methods

### Description of RNN and training

An RNN is composed of *N* artificial neurons (or units) that receive input from *N*_in_ time-varying inputs **u**(*t*) and produce *N*_out_ time-varying outputs **z**(*t*). The RNN defines a network state, given by *x*(*t*) ∈ ℝ^*N*^; the ith element of **x**(*t*) is a scalar describing the “currents” of the *i*th artificial neuron. The network state is transformed into the artificial neuron firing rates (or network rates) through the transformation:

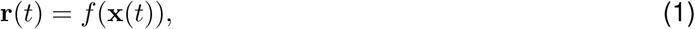

where *f*(·) is an activation function applied elementwise to x(*t*). The activation function is typically nonlinear, endowing the RNN with nonlinear dynamics and expressive power. In this work, we use *f*(*x*) = tanh(*x*) as well as *f*(*x*) = max(*x*,0), also known as the rectified linear unit or relu(·). In the absence of noise, the continuous time RNN is described by the equation

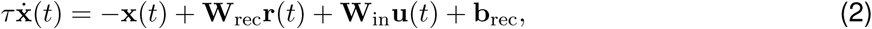

where *τ* is a time-constant of the network, **W**_rec_ ∈ ℝ^*N*×*N*^ defines how the artificial neurons are currently connected, b_rec_ ∈ ℝ^*N*^ defines a constant bias, and **W**_in_ ∈ ℝ^*N*^×*N*_*in*_ maps the RNN inputs onto each neuron. We note that equation 1 can also be used to calculate the dynamics of the network rates, **ṙ**(*t*). This quantity is useful because it describes how the network rates evolve through time. In neurophysiological studies, this is equivalent to calculating the dynamics of the recorded neuron firing rates (Churchland et al., 2012; Kao et al., 2015).

The output of the network is given by a linear readout of the network rates, i.e.,

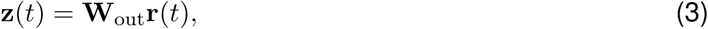

where **W**_out_ ∈ ℝ^*N*_out_×*N*^ maps the network rates onto the network outputs. We trained the RNN to minimize the mean-square error between its output, **z**(*t*), and a desired output, **z**_des_(*t*). In addition to this objective, we included several regularizations to improve training. In particular, we regularized the *L*_2_-norm (Euclidean norm) of **W**_in_, **W**_rec_, and **W**_out_ to penalize larger weights. We also regularized the *L*_2_-norm of **r**(*t*) across all time, as was done in (Sussillo et al., 2015; Michaels et al., 2016) to penalize larger rates, which are not encountered in biological neurons due to their refractory period. We later report that this regularization has an important impact on firing rates during movement preparation. Finally, we incorporated gradient clipping and the regularization proposed by Pascanu and colleagues to ameliorate vanishing gradients (Pascanu et al., 2012). Training was performed using stochastic gradient descent, with gradients calculated using backpropagation through time. For gradient descent, we used the Adam optimizer, which is a first order optimizer incorporating adaptive gradients and momentum (Kingma & Ba, 2014). Finally, because reaching behavior is highly stereotyped, we allowed training to continue until the coefficient of determination in kinematic reconstruction on validation data exceeded *R*^2^ × 0.997.

### Visualizing RNN dynamics

The RNN’s dynamics are fully described by equation 2. Thus, one may qualitatively assess the RNN’s dynamical mechanism by visualizing this equation. However, in most scenarios, the RNN is composed of a relatively large number of neurons, *N* (e.g., typically *N* > 100). By treating each artificial neuron as an independent dimension, this implies that **ṙ**(*t*) is *N*-dimensional, and thus not trivial to visualize. One way to address this problem is to consider that, in many scenarios, not unlike what is observed in neural population activity in motor cortex (Yu et al., 2009; Cunningham & Yu, 2014), the dimensionality of the *N* artificial neurons are correlated and can thus be adequately described in a *D* dimensional subspace, where *D* < *N*. Hence, while the dynamics implemented by the RNN cannot be visualized in a straightforward manner if *N* is large, it may be possible to do so if the dynamics can be appropriately sampled in *D* = 1 to 3 dimensions.

There are important considerations in visualizing dynamics in a low-dimensional subspace. The primary consideration is that the dynamics in any *D*-dimensional projection will differ depending on the activity in the remaining *N – D* dimensions. Hence, depending on the values the network rates take on in the remaining *N – D* dimensions, the visualized dynamics may differ in a minor or substantial way. To illustrate this concept, consider the 3D example shown in Fig 1a. As shown in Fig 1a, we can measure the firing rates of each artificial neuron for a given input, and plot the trajectory of **r**(*t*) in 3 dimensions, where each dimension is defined by the activity of one artificial neuron.

In Fig 1e, we introduce the notion of projecting the RNN dynamics into a given subspace. Consider an orthonormal basis given by **U**_3_ = [**u**_1_ **u**_2_ **u**_3_] where each **u**_*i*_ ∈ ℝ^*N*^ and 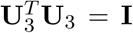. We can define a two-dimensional trajectory by projecting the network rates into the subspace spanned by **U**_2_ = [**u**_1_ **u**_2_]. The low-dimensional trajectory in this subspace is given by

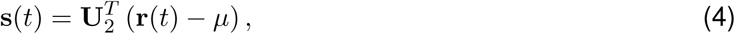

where *μ* is the mean of **r**(*t*) across time, and its dynamics can be calculated as

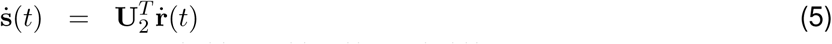

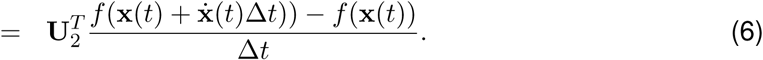

However, this sampling rule is naïve in the following way. In Fig 1e, we consider the trajectory at times *t*_1_ and *t*_2_, denoted by **r**(*t*_1_) in red and **r**(*t*_2_) in green, respectively. If the trajectory **r**(*t*) has a relatively large component along **u**_3_, the dynamics may be very different (e.g., at time *t*_2_, illustrated by the green plane). This is because:

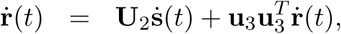

and thus, the low-dimensional dynamics embedded in the high-dimensional space (given by **U**_2_ṡ(*t*), and plotted as the flow field trajectories in Fig 1e) may change if 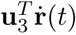 is large. We note that it is possible to sample the dynamics accurately at any single time point *t*, since the component of **r**(*t*) in the orthogonal complement of **U**_*D*_ is known. Hence, it is possible to visualize local dynamics over time in a movie by re-sampling the low-dimensional dynamics at every time t for a given **U**_*D*_; such a movie is shown in Supplementary Movie 1.

To address the changing dynamics, we propose two heuristics to find the low-dimensional subspace, **U**_*D*_ (where *D* is the number of dimensions) to sample RNN dynamics. We wish to find a matrix **U**_*D*_ with *D* orthonormal columns so that 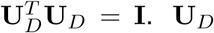. **U**_*D*_ defines the subspace where we will visualize the network rates **r**(*t*) as well as their dynamics **ṙ**(*t*). With this definition, 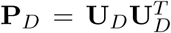 is a projector matrix into the subspace spanned by **U**_*D*_. To create a meaningful dynamical portrait from which it may be possible to glean intuition as to how the RNN performs a given task, the subspace should capture meaningful variance in the data, as *well as* capture a faithful view of the dynamics. We enumerate two heuristics to sample these dynamics:

1. Intuitively, the components of **r**(*t*) along the remaining *N* – *D* dimensions do not change dramatically. In this manner, the sampled D-dimensional subspace is approximately the same across time. In the context of Fig 1f, this corresponds to the red and green subspaces being relatively close to each other. Assuming a smoothness in the RNN dynamics, if this separation is sufficiently small, the dynamics will not change drastically. This smoothness assumption is valid for the tanh nonlinearity, but for the relu nonlinearity fails at x(*t*) = 0. This projection has the added benefit of finding the projection that maximizes the projected data variability 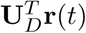. Formally, this projection can be found by maximizing the variance of 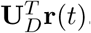. The solution of this optimization is called principal components analysis, where **U**_*D*_ correspond to the first *D* left singular vectors of [**r**(1) **r**(2) … **r**(*T*)], with T being the horizon of the data.
2. Intuitively, the projected dynamics in “dynamics relevant” dimensions ought to be oriented in similar directions over the course of the epoch. This reflects that the dynamics will not vary substantially over the course of the epoch, as illustrated in Fig 1g. This may be achieved by optimization of an appropriate loss function over Stiefel manifolds.

In this work, we found that heuristic 1, projection along principal components, was sufficient for our analyses. While we also performed an optimization under heuristic 2, we found that this did not affect any conclusions. After finding **U**_*D*_, we visualized the network rates and their dynamics by using equations 4 and 5 (and substituting *U*_*D*_ for **U**_2_), respectively. Finally, we also visualized the fixed points of the RNN’s dynamics using the approach presented by Sussillo and Barak (Sussillo & Barak, 2013).

### RNN task and training

We trained the RNN on a variant of the delay task used by Ames and colleagues (Ames et al., 2014). This study proposed a specific dynamical mechanism that we sought to probe with RNNs. In this task, a monkey is instructed to hold a center target. After holding the center target for 700 – 1100 ms, a peripheral target is cued in one of eight locations uniformly spaced on a circle, 45° apart beginning at 0°. The monkey continues to hold the center target while planning to reach to the cued target. After a random delay period, ranging from 0 – 900 ms, the monkey is given a go cue and is allowed to perform a reach to the prompted target. Upon reaching the target, the monkey then holds on the target for a 500 ms hold time to successfully acquire the target, ending the trial. This task is diagrammed in Fig 2a.

**Figure 2:**
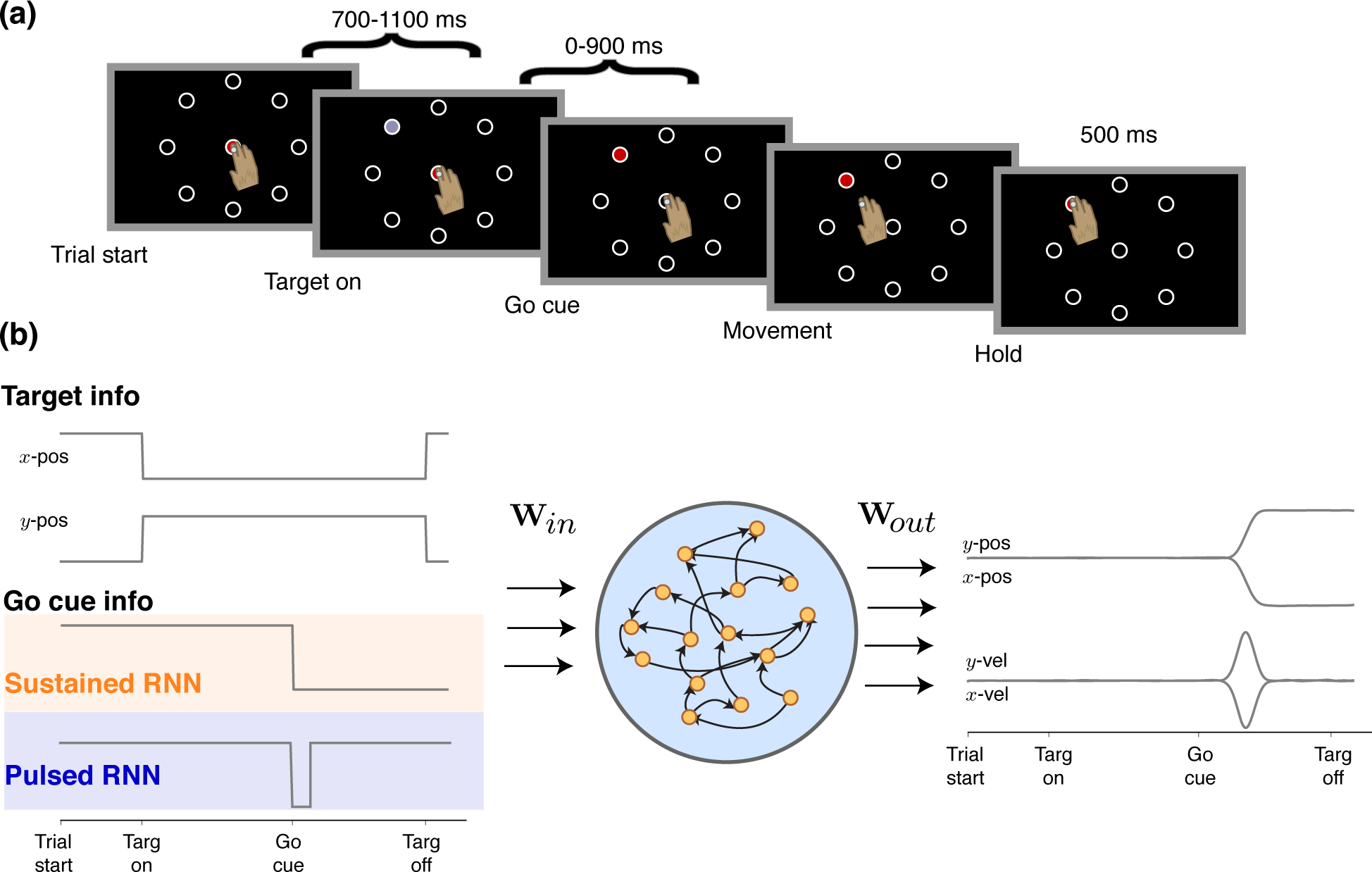
Delayed reach task and RNN training. **(a)** Schematic of the RNN task used by Ames and colleagues. For RNN training, the hold time was allowed to be variable for anywhere from 500 to 1500 ms so that the RNN did not learn specific timings. **(b)** Example (and representative) target inputs and outputs of the RNN. The RNN was trained until it achieved a coefficient of determination, *R*^*2*^ > 0.997, on the task. The RNN outputs both position and the velocity. The sustained RNN encodes the go cue as a signal to withhold movement (1) or to allow movement (0) while the pulsed RNN encodes the go cue as a transient cue that indicates a change in the state of the task (i.e., that the monkey can now execute his planned reach).

The RNN inputs were the target’s *x*-position, the target’s *y*-position, and a go cue signal. Because we were interested in assessing the effect of inputs on dynamical mechanisms, we trained with two different go cue configurations. In the “sustained RNN” (Fig 2b, orange) the go cue was encoded with a sustained signal indicating whether to withhold movement (signal high) or not (signal low) as in prior studies (Sussillo et al., 2015; Michaels et al., 2016). In the “pulsed RNN” (Fig 2b, blue) the go cue was encoded as a transient pulse indicating that movement could occur. This go pulse could correspond to when the animal is cued that he may move by a transient cue (e.g., a brief and temporary visual cue). This pulse may also be interpreted as reflecting that the state of the task has changed, so that the animal may now make a reach, analogous to a signal that triggers movement (Erlhagen & Schöner, 2002; Kaufman et al., 2016). An additional motivation for using the pulse is that prior tasks have used transient cues, such as networks trained to process a transient movement instruction (Hennequin et al., 2014). The RNN transformed these inputs into four outputs: the *x*‐ and *y*-positions of the hand, and the *x*‐ and *y*-velocities of the hand. In this manner, the input was 3-dimensional, **u**(*t*) ∈ ℝ^3^, and the output was 4-dimensional, **z**(*t*) ∈ ℝ^4^. Our trained networks had 100 artificial neurons, so that x(*t*), **r**(*t*) ∈ ℝ^100^.

Like in the study by Ames and colleagues, we trained the network with reaches having delay periods ranging from 0 – 900 ms after a 700 – 1100 ms center-hold period. After each delay period, we had the network produce a reach following a fixed reaction time of 150 ms. After the reach transient, the network was then trained to generate zero *x*‐ and *y*-velocities and appropriate final positions for a hold period. Instead of a static 500 ms hold period used by Ames and colleagues, we allowed the hold period to be from any length from 500 to 1500 ms, so that the network didn’t learn specific timings (i.e., to use a region of slow dynamics for only 500 ms). We trained the network to produce reaches to 8 targets uniformly spaced on a circle. The targets were 45° apart beginning at 0°. In addition to these delayed reaches, on 10% of trials, we introduced “catch trials” to the RNN where a target may not have appeared, or if it appeared, the go cue was never given. In both instances, the network had to sustain zero output.

In the task by Ames and colleagues, there were also occasional *switch* trials, where the target was switched on 20% of trials. Following this target switch, the monkey was given a second delay period ranging from 0 – 900 ms followed before the go cue was delivered. We explicitly did not train on this task because we were interested in assessing how the RNN would generalize to it.

## Results

Before delving into design choices, we found that it was possible to train an RNN to recapitulate key features of the neural activity in a delayed reach task, as reported in prior studies (Sussillo et al., 2015; Michaels et al., 2016). The hyperparameters for this network are listed in Supplementary Table 1. Fig 3a shows peristimulus time histograms (PSTHs) of artificial neurons for delayed reaches to eight different directions. These PSTHs plateau during the delay period (Tanji & Evarts, 1976; Weinrich et al., 1984; Churchland et al., 2010; Sussillo et al., 2015; Michaels et al., 2016), have substantial heterogeneity and multiphasic activity during perimovement (Sergio et al., 2005; Churchland & Shenoy, 2007; Churchland et al., 2010), and change preferred directions over time (Churchland & Shenoy, 2007; Michaels et al., 2016), reflected by the fact that the condition with the highest firing rate is not the same across the entire trial. Fig 3b-e shows that the artificial neural population also recapitulates qualitative observations from neurophysiologically recorded neural populations. We found that only 5 PCs were required to capture more than 90% of the PSTH variability, as shown in Fig 3b, demonstrating that the population is low-dimensional (Yu et al., 2009; Cunningham & Yu, 2014; Ames et al., 2014; Sadtler et al., 2014; Kao et al., 2015; Gallego et al., 2017; Gao & Ganguli, 2015), that PC_1_, capturing 43.7% of the PSTH variability, strongly resembled a high variance condition-independent signal (Kaufman et al., 2016) (Fig 3c), that artificial neural population activity had topographic organization in the PCs (Santhanam et al., 2009) (Fig 3d), and that the neural population achieved a prepare-and-hold state attractor (Churchland et al., 2006) (Fig 3d) but that this attractor was not obligatory (Ames et al., 2014) (Fig 3e).

**Figure 3:**
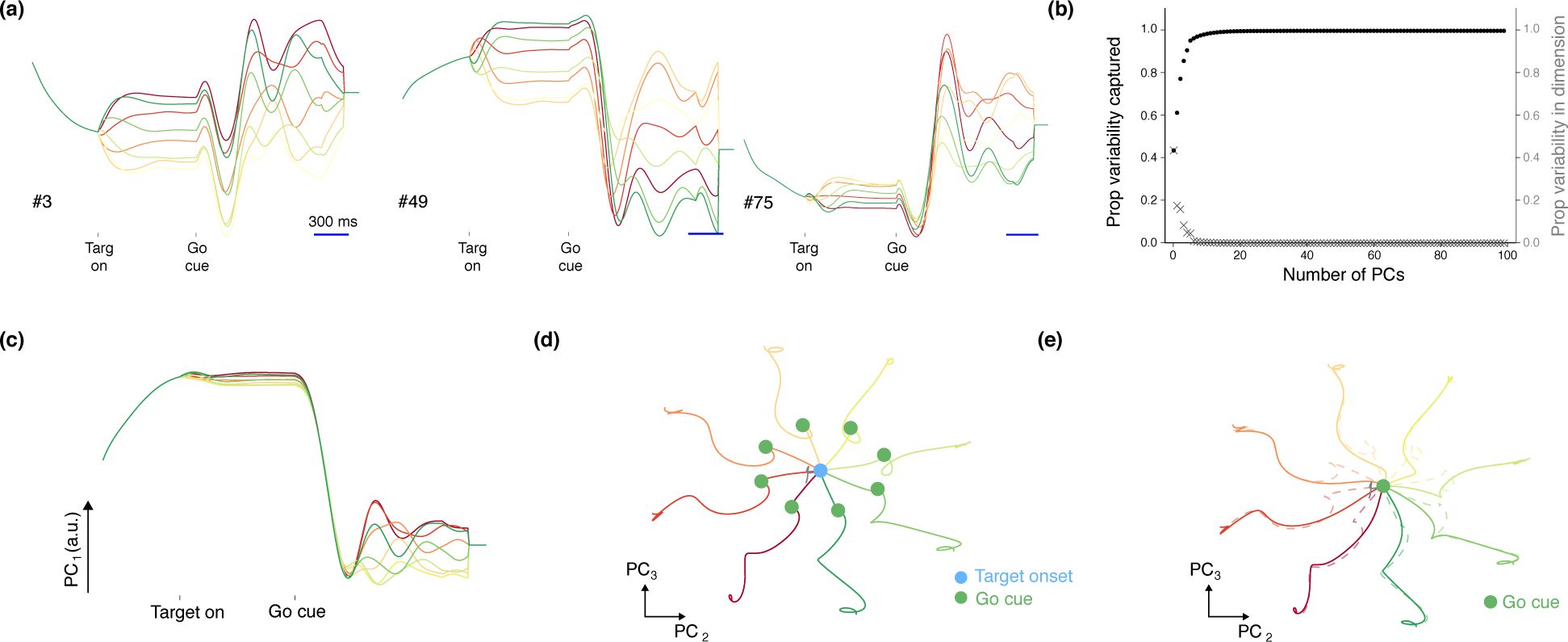
Sample PSTHs and population trajectories of the RNN. **(a)** Example PSTHs of artificial neurons in the RNN, where each color denotes one of the eight possible reach conditions. The PSTHs achieve a stable firing rate in the preparatory period, followed by multiphasic activity during perimovement. The condition with the highest firing rate may also change through time. **(b)** Proportion of variance captured by the principal components. 5 PCs capture 90.8% of the PSTH variability, and 10 PCs 98.2% of the variability. **(c)** PCi, capturing 43.7% of the signal variance, demonstrates properties consistent with the condition independent signal proposed by Kaufman and colleagues. It is the largest response component of the RNN rates, but largely does not reflect movement type (until well after the go cue has been delivered). **(d)** RNN rate trajectories in the PC_2_ and PC_3_ axis with a delay period. For each reach condition, the trajectories reach a stable set of neural states prior to go cue delivery. These are indicated by the trajectory locations at the green dot. Adjacent conditions are topographically organized (0°: red, 45°: dark orange, 90°: orange, 135°: light orange, 180°: yellow, 225°: light green, 270°: green, 315°: dark green). The gray part of the trajectory represents the baseline activity. **(e)** Trajectories (in bold) when the RNN performs the task without a delay period. This shows that the preparatory neural states are not obligatory, consistent with the findings of Ames and colleagues.

### Rate regularization and activation function affect preparatory activity

Prior studies using RNNs to model motor cortex use the hyperbolic tangent (tanh) activation function (Michaels et al., 2016; Sussillo et al., 2015) or a variant of it (Hennequin et al., 2014). Recent studies have also used the rectified linear unit (relu) nonlinearity to model various decision making tasks (Song et al., 2016, 2017). We note the relu nonlinearity has proliferated in several engineering applications, in part due to the faster training times and that the gradient of the relu is either zero or one, which is favorable for backpropagation (Krizhevsky et al., 2012; Szegedy et al., 2015; He et al., 2016). We found that the choice of activation function impacts preparatory activity during the delay period. Preparatory activity captures a substantial proportion of neural variability. Typically, preparatory activity plateaus to a stable level prior to movement onset (Tanji & Evarts, 1976; Weinrich et al., 1984; Churchland et al., 2010; Sussillo et al., 2015; Michaels et al., 2016). In a dynamical systems framework, the population preparatory activity evolves to a subspace called the “prepare-and-hold” state that is beneficial for the upcoming reach (Churchland et al., 2006; Afshar et al., 2011; Ames et al., 2014).

Given its prior use in RNN models of motor cortex, we first considered the hyperbolic tangent nonlinearity. Interestingly, we found that rate regularization (weighted by λ_*r*_) was important for achieving preparatory activity that was qualitatively consistent with neurophysiological data. When rate regularization was relatively small, we found that artificial neurons in the RNN had little preparatory activity (Fig 4a, leftmost panel). This can be observed by recognizing that population activity at the time of the go cue essentially overlapped with population activity at target onset. This solution is not unreasonable because the RNN’s outputs remain zero during both the baseline and preparatory epochs. While target information is available to the RNN in the preparatory period, it does not necessarily have to act (i.e., change its state) upon this information until the go cue is given. To this end, the RNN can delay processing target information until the go cue is given and still successfully perform the task.

**Figure 4:**
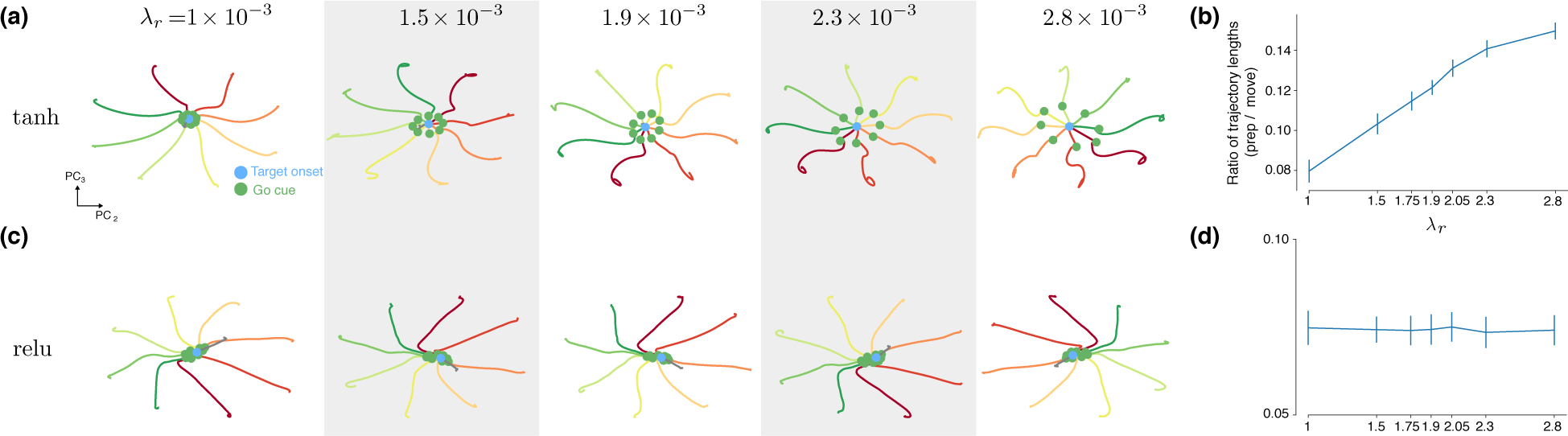
Rate regularization increases preparatory activity in tanh but not relu RNNs. **(a)** Population trajectories of tanh RNNs in PCs 2 and 3 (analogous to Fig 3d) for different values of λ_*r*_, which weights the *L*_2_ norm penalty on the rates. As λ_*r*_ increases, the preparatory trajectory becomes larger relative to the movement trajectory. **(b)** Ratio of the the preparatory trajectory length (in all N dimensions, not only in selected PCs) divided by the movement trajectory length. As λ_*r*_ increases, the preparatory trajectory length becomes relatively larger. Error bars are standard deviations across 8 separate RNNs trained at each value of λ_*r*_. **(c)** Same as **(a)** but for relu. As λ_*r*_ increases, there is not a noticeable increase in preparatory activity. **(d)** Same as **(b)** but for relu.

We found that, for the hyperbolic tangent nonlinearity, increasing rate regularization increased the amount of preparatory activity in the network. This is shown for several values of λ_*r*_ in the remaining panels of Fig 4a, and is summarized by Fig 4b, which shows the ratio of lengths between the preparatory trajectory and the movement trajectory. The trajectory ratios are calculated in the high-dimensional artificial neuron activity space and not in the low-dimensional PCs. By observing the PSTHs of the activity at different levels of regularization (Supp Fig 1), we observe that rate regularization causes the rates to achieve (1) smaller overall peak values and (2) intermediate activations in the preparatory epoch. In this manner, rate regularization causes the tanh RNN to have stronger preparatory dynamics, effectively partitioning computation into two segments: a preparatory dynamical system (driving the activity to a fixed point, denoted by the green dots in Fig 4a) followed by the movement dynamical system (trajectory after the green dot). This is most apparent in the rightmost panel of Fig 4a. We note that this population activity is consistent with what is qualitatively observed in motor cortex. For the hyperbolic tangent nonlinearity, this is an energetically favorable outcome for the network.

We emphasize that increasing rate regularization does not always result in more preparatory activity. In fact, when we used the relu activation, we observed empirically that the network finds a solution that has little preparatory activity irrespective of λ_*r*_ (Fig 4c; trajectory lengths summarized in Fig 4d). We note that this was not because rate regularization was not “active” due to other regularizations dominating the optimization cost; in fact, when we removed all regularization except rate regularization, the networks still had units with very little preparatory activity across 4 orders of magnitude (from λ_*r*_ = 10^‒3^ to λ_*r*_ = 10). PSTHs of the relu RNN are shown in Supp Fig 2, and demonstrate that although the maximum rate may decrease as rate regularization increases, the preparatory activity does not appear to increase in variability relative to movement activity. Togther, these results suggest that the relu() activation, for these instantiations of RNNs, does a poorer job of capturing key features in the neurophysiological activity. Further, we point out that it was not the case that units with preparatory activity were absent in the network. Rather, Supp Fig 2 demonstrates that it was possible to find relu units that had substantial preparatory activity. Hence, when considering RNNs to model neurophysiological tasks, its important to not only find single unit examples that resemble physiological activity, but that also recapitulate population level features. For the rest of this work, we utilize the tanh() activation with rate regularization λ_*r*_ = 1.9 × 10^‒3^.

### RNN dynamics in the sustained RNN during a delayed reach task

We next visualized the dynamics of the RNN (see Materials and Methods) as displayed in Fig 5. We also visualized the stable attractor regions of the dynamics by using the approach of (Sussillo & Barak, 2013). We found that the RNN implemented a dynamical mechanism that can be interpreted as a composition of dynamical systems to single stable attractor regions. Key features of this mechanism were proposed by Ames and colleagues to describe neural population activity during a delayed reach task (Ames et al., 2014). In particular, Ames and colleagues proposed two principal dynamical systems: a “preparatory” dynamical system implicated in planning a reach to a desired target, and a “movement” dynamical system corresponding to the execution of the reach after the go cue is delivered. The preparatory dynamical system has a stable attractor corresponding to each prompted target. The neural state converges to this attractor, the prepare-and-hold state, during the delay period. This state is a favorable initial condition for the subsequent movement (Churchland et al., 2006; Afshar et al., 2011). When the go cue is given, the movement dynamical system is engaged, driving the trajectory through a path associated with movement generation.

**Figure 5:**
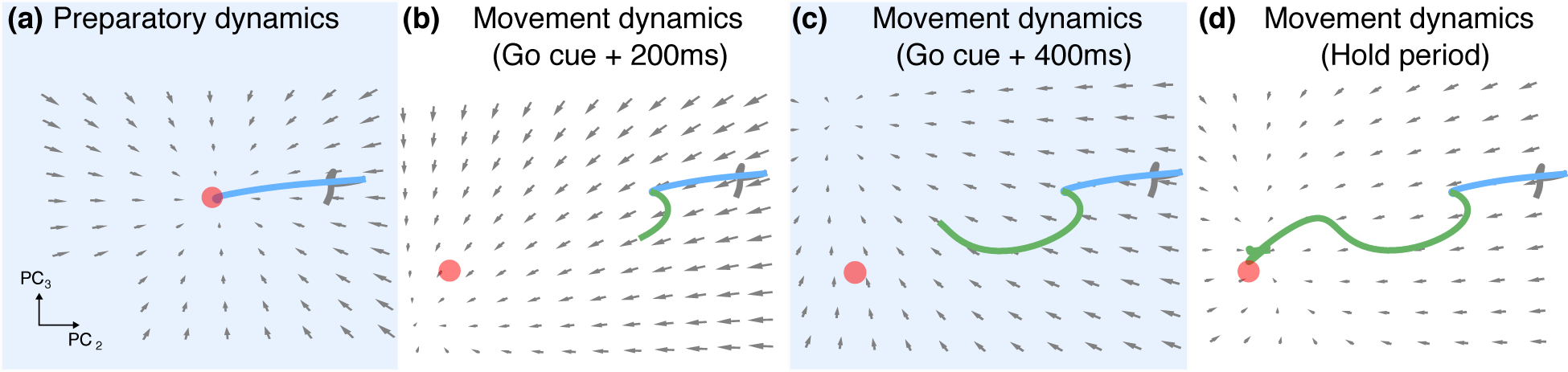
Composition of dynamical systems mechanism. **(a)** The delay period trajectory in PCs 2 and 3. The gray portion of the trajectory corresponds to the baseline period of the task. The blue portion of the trajectory corersponds to the delay period of the task. The state converges along essentially linear dynamics to a stable attractor (shown as a red dot). The attractor was found as the location in state space that minimized ||ẋ||^2^ below a threshold tolerance. **(b)** The movement dynamics 200 ms after go cue onset. The green portion of the trajectory corresponds to the post go cue period of the task. The dynamics are strongly driven towards a single stable attractor region. There appear to be non-zero dynamics overlapping the attractor because the trajectory is not in the same plane as the attractor (in orthogonal dimensions). **(c)** The movement dynamics 400 ms after go cue onset. The dynamics have changed due to trajectory movement in the orthogonal dimension. **(d)** Approaching the hold period, we see that the dynamics converge on the stable attractor.

By visualizing the RNN’s dynamical equations, we were able to qualitatively probe how the RNN uses nonlinear dynamics to perform the task. We found that during the delay period, the RNN implemented an analogous preparatory dynamical system. Upon target presentation, the trajectories were driven by this preparatory dynamical system to a single stable attractor region as in Fig 5a. This stable attractor location was target dependent. The RNN achieved different preparatory attractors and dynamics for a given target because the network’s dynamics at a given time, ẋ(*t*), are modified by an input-dependent additive factor, **W**_in_**u**(*t*). Hence, each unique input can be interpreted as setting up a unique dynamical system. This enables the network to, prior to the go cue, instantiate different preparatory dynamics with different stable attractor regions for each prompted target.

When the go cue was delivered, the changing input drastically modified the dynamics, so that the trajectories were driven along paths associated with output generation (i.e., the movement dynamical system). We found that this movement dynamical system was comprised of a single stable attractor region. The movement dynamical system utilized strong dynamics to drive the RNN’s rate trajectories at relatively high speeds, as shown in Fig 5b,c. In this manner, the mechanism is not integration along slow points on a line attractor (e.g., Mante et al. (2013)), but rather abrupt state transitions from stable attractor to stable attractor. A video of these dynamics is shown in Supp Movie 1. These types of dynamics, illustrated in Fig 1c, have been observed in another study (Chaisangmongkon et al., 2017). An increase in the speed of neural trajectories following the go cue is consistent with experimental observations from PMd and M1 (Afshar et al., 2011) (their Fig 3c). Note that when target presentation is simultaneous with the go cue so that there is no delay period, the movement dynamical system is immediately engaged, and trajectories are driven by the movement dynamical system to its single stable attractor region. As a result, the preparatory dynamical system attractor is not obligatory. Because the preparatory dynamical system has not been engaged for enough time, the trajectories will not achieve the preparatory attractor, a phenomena also observed in neurophysiological data (Ames et al., 2014).

### RNNs qualitatively recapitulating neurophysiological motifs may utilize different dynamical mechanisms

We next wondered if task design considerations could produce RNNs that, while looking qualitatively similar to neurophysiological data, utilize distinct dynamical mechanisms. To this end, we trained the earlier described pulsed RNN to perform a delayed reach task, using the same hyperparameters as the sustained RNN. In the training set, the pulsed go cue was delivered for 150 ms. This pulse may also be interpreted as reflecting that the state of the task has changed, so that the animal may now make a reach, analogous to a signal that triggers movement (Erlhagen & Schöner, 2002; Kaufman et al., 2016). We are not suggesting that this pulse length would be reasonable for experiments; although we chose to use 150 ms, this pulse length can be varied. The RNN was capable of performing the pulsed go cue task at the same level as the single attractor RNN (training terminated when *R*^2^ > 0.997 on validation data, example output trajectory shown in dark blue in Fig 6a). Its recurrent computation was similarly low-dimensional, with 5 PCs capturing 91.8% of the PSTH variability (Supp Fig 3a). This RNN also bore hallmarks of neurophysiological responses, including: neural activity being organized topographically (Supp Fig 3f), the trajectories achieving a “prepare-and-hold” state in the delay period (Supp Fig 3f), and that these states were not obligatory (Supp Fig 3e). We do note that condition-independent variance, though present, appeared to be smaller in this network, with a large proportion appearing in PC 3 (Supp Fig 3b-d).

**Figure 6:**
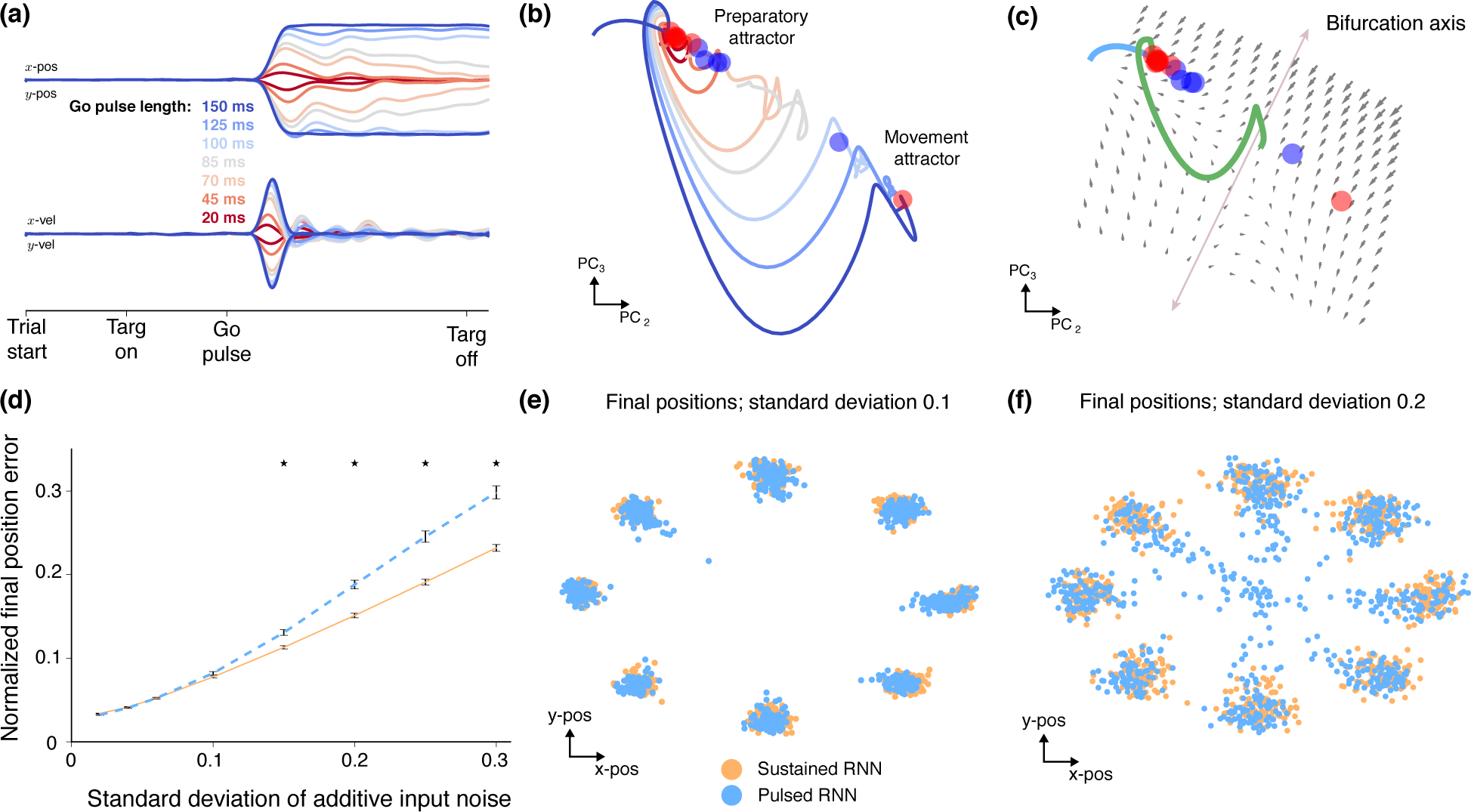
An RNN trained to perform a delayed reach task with a transient go cue. **(a)** Example output for an RNN that was trained on a pulsed go cue task to make a reach to the target at 315°. The output is shown for pulsing the go cue at different lengths, denoted by different colors. When the go cue is pulsed for greater than 85 ms, the RNN eventually outputs the correct final *x* and *y*-positions. When the go cue is pulsed for 85 ms or less, the RNN decays to output incorrect final zero *x*‐ and *y*-positions. **(b)** Neural trajectories for the reach to the target at 315° for different length go pulses. The trajectories either decay back to the preparatory state (left attractor region) or eventually converge to the stable attractor associated with movement generation (right attractor region). Red circles denote stable slow regions of state space; blue circles denote unstable slow regions of state space. **(c)** The dynamics at the point of bifurcation. The bifurcation axis is illustrated in light purple. Left of the axis, the dynamics will drive the trajectory to decay back to the preparatory attractor. Right of the axis, the dynamics will eventually drive the trajectory to the attractor associated with the correct final output. **(d)** The y-axis denotes the normalized final position error (normalized so that the final position is 1). The x-axis denotes the standard deviation of independent zero mean Gaussian noise added to the inputs. The dotted line represents the performance of the pulsed RNN, while the solid line represents the performance of the sustained RNN. As input noise increases, the pulsed RNN has worse final position performance. Stars denote significant differences in the mean (bootstrap, 1001 shuffles, *p* < 0.01). **(e)** Final positions to the eight targets for RNNs of both mechanisms when the input noise standard deviation is 0.1. Each dot represents the final position on a single trial. Both RNNs still generate relatively accurate outputs. **(f)** When the input noise standard deviation is increased to 0.2, the pulsed RNN has several trials where the final position begins to decay back to the center target, which is the kinematic output corresponding to the preparatory attractor. The hold time was increased to 2000 ms to show this slow decay. Trials which end at intermediate locations may reflect trajectories in slow regions of decay back to the preparatory attractor, as well as variable end points due to large input noise.

In this task design, the input during the delay period is the same as the input during the movement period post go cue. We reasoned that under the insight that each input changes the RNN’s dynamics, as seen in Fig 5 and Supp Movie 1, this RNN cannot use a composition of dynamical systems with single stable attractor regions. Doing so would imply that the delay and movement periods must converge to the same stable attractor, and hence the delay and movement periods would converge to the same output under this mechanism. Such an RNN would be unable to adequately perform this task. We note that while we have chosen a pulsed go cue, a similar conclusion holds for RNNs which use transient cues, such as the networks trained by Hennequin and colleagues with a movement instruction cue (Hennequin et al., 2014). Analogously, this network produced two different output transients (pre-preparatory and post-movement) for the same input.

Although the trajectories in condition-relevant dimensions demonstrate qualitatively similar trajectories to neurophysiological data, how does the RNN dynamically achieve this, if not by the mechanism used by the sustained RNN? To answer this question, and recognizing the RNN must be capable of achieving two steady-state outputs, we pulsed the go cue to determine what duration of go cue was required for the pulsed RNN to settle to the correct final kinematics as opposed to returning to the preparatory state kinematics (i.e., zero positions and velocities). We delivered different input pulses, as shown in Fig 5a, and found that when the pulsed go cue was delivered for less than 85 ms, the pulsed RNN produced transient kinematics that decayed back to zero. However, when the go cue was delivered for greater than 85 ms, the pulsed RNN produced the correct final kinematic output corresponding to the prompted reach.

By visualizing the trajectories and dynamics, as shown in Fig 6b,c, it is clear that the pulsed RNN sets up at least two stable attractor regions. Our optimization did not identify any other attractors, suggesting that the pulsed RNN implements a bistable dynamical system. One stable state is associated with the preparatory period, analogous to the prior RNN, where the RNN rates converge to during the delay period. The second stable state is the region associated with making a movement to the prompted target. By visualizing the dynamics, shown in Fig 6c, we were able to identify bifurcation dynamics associated with the task. These bifurcation dynamics could be viewed in the projections of PCs 2 and 3. The go cue pulse drives the trajectory to this region of bifurcation, and depending on the trajectory’s location, it will either settle back to the preparatory state (with zero kinematic output, i.e., left of the illustrated bifurcation axis) or settle to the state associated with the correct kinematic output (i.e., right of the illustrated bifurcation axis).

These results demonstrate that, for trajectories having qualitative similarities to neurophysiologically observed data, different dynamics may be at play. While both RNNs utilize a composition of dynamical systems (Ames et al., 2014), we found that the employed mechanism differed substantially from task input design (i.e., strongly driving trajectories to single attractors versus implementing a region of bifurcation). In considering how to then militate between these two mechanisms, we asked which mechanism was more robust to input noise, as could occur from suboptimal processing of the task inputs. To this end, we added independent zero-mean Gaussian noise to the inputs, and assessed the RNN’s performance as a function of the standard deviation of the Gaussian noise. We found that increasing the input noise affects the network in at least two distinct ways.

First, for both RNNs, because the stable attractor region is input-dependent, noisier inputs cause the stable attractor region to be variable, resulting in greater neural trajectory end point variability, and hence, kinematic end point variability. We observed this effect, as end point deviation increased with the standard deviation of Gaussian noise (Fig 6d) and in Fig 6e, substantial variability can be observed in the end positions. Interestingly, however, we did not observe a significant difference in performance between the sustained and pulsed RNNs when the standard deviation was less than or equal to *σ* = 0.1, which is approximately 10% of the input signal (*p* = 0.69, bootstrap with 1001 shuffles). Hence, the effect of input noise varying attractor location was similar in both networks.

Second, we found that for the pulsed RNN, input noise may cause the trajectory to not cross the bifurcation axis, resulting in eventual relaxation to the preparatory attractor. As shown in Fig 6d, when the standard deviation increased beyond *σ* = 0.15, the pulsed RNN had worse final end point performance than the single attractor RNN (*p* < 0.01, bootstrap with 1001 shuffles). Importantly, for the pulsed RNN, we observed that its state occasionally relaxed back to the preparatory attractor, corresponding to a (0,0) output center position, as can be observed by the final end point of the kinematics shown in Fig 6f. This demonstrates that in the presence of noise, RNN task performance will similarly degrade until the point where noise causes the pulsed RNN’s state to converge to the incorrect attractor. This suggests that, for the purposes of performing a delayed reach task, the sustained RNN is more robust under input noise.

### RNN generalization to new tasks

Finally, we wondered the extent to which a qualitative understanding of the RNN’s dynamics could inform task generalization. We believe this is an important line of questioning for future RNN studies. In particular, by constraining what tasks the RNN is trained on, it is possible to comment on what dynamical mechanisms are sufficient to carry out certain tasks. As an example, can an RNN using the composition of preparatory and movement dynamical systems generalize to perform a target switch task? The target switch task is diagrammed in Fig 7a. We hypothesized this ought be possible; indeed, Ames and colleagues used a qualitatively consistent dynamical mechanism to describe how motor cortex performs a target switch task. By training an RNN to only perform a delayed reach task and utilizing the composition of dynamical systems, we can assess whether this dynamical mechanism is sufficient to enable the network to perform related tasks it was not trained on. We considered three variants of a target switch task, where the target switches at different times: (1) before the go cue, (2) simultaneous with the go cue, and (3) after the go cue. The first two were considered in neurophysiological experiments by Ames and colleagues (Ames et al., 2014).

**Figure 7:**
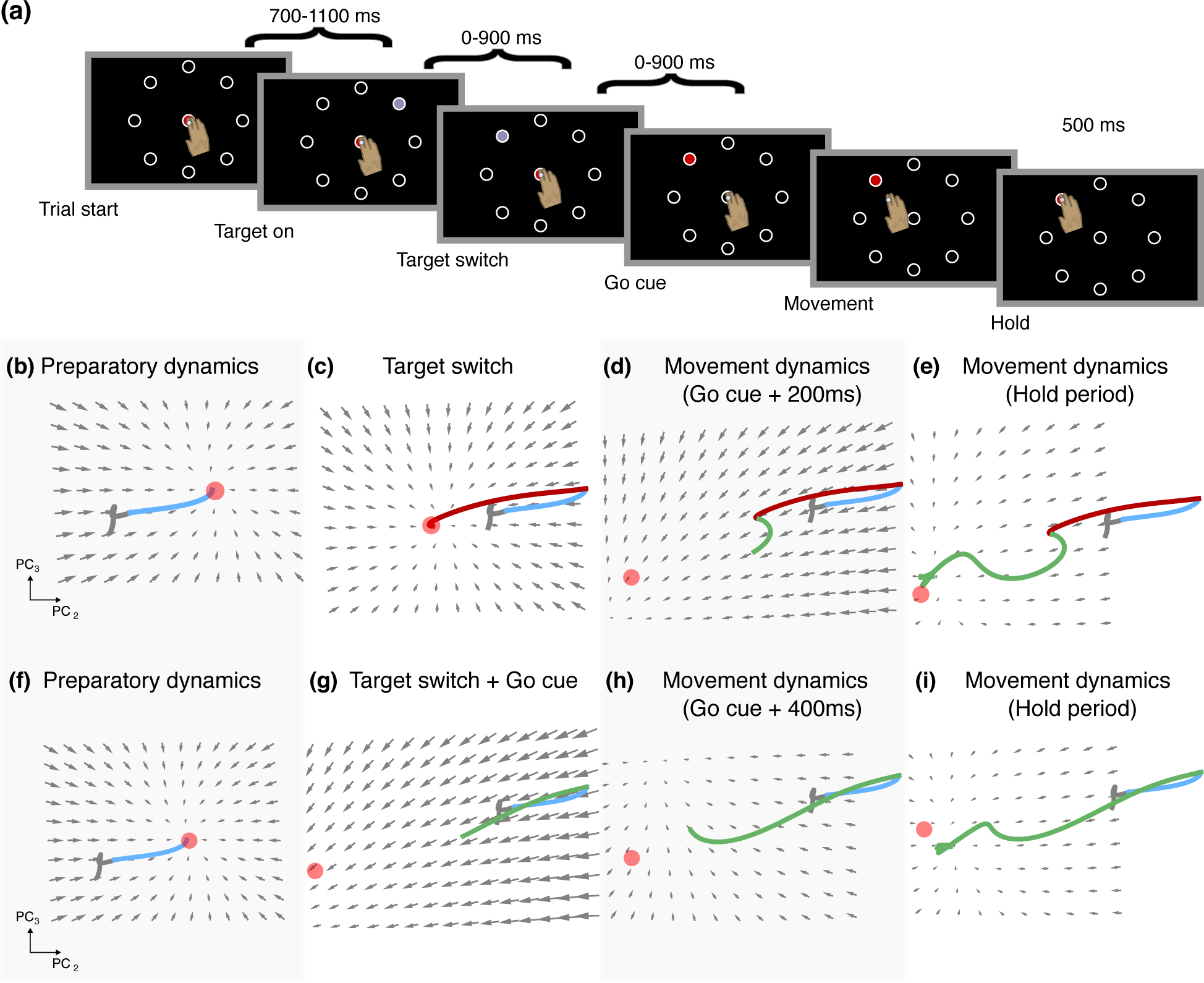
RNN dynamics during a target switch task. **(a)** Schematic of the target switch task when the go cue is delivered either with a delay or contemporaneously with the go cue. **(b-e)** RNN dynamics during a target switch task, with an additional delay period to re-prepare. **(b)** Preparatory dynamics during the delay period (blue trajectory). **(c)** Dynamics during the switch period (red trajectory). The new target changes the stable attractor region, and the dynamics drive the trajectory to this attractor. **(d-e)** Dynamics following the go cue, recapitulating the same trajectory seen in Fig 5c,d. **(f-i)** Same as **(b-e)** but when the go cue is given contemporaneously with the target switch.

Consider first the sustained RNN. When the target switch is delivered prior to the go cue, the preparatory dynamical system is changed from that associated with the before-switch target to that of the after-switch target. As a result, the preparatory stable attractor changes when the target is switched, and the RNN’s rates will converge to the single stable attractor associatd with the switched target. When the go cue is then delivered, the RNN will execute the reach as it did in a delayed reach task. We found this was the case, as illustrated in Fig 7b-e, mimicking the experimental results presented by Ames and colleagues (Ames et al., 2014). When the go cue is given simultaneously with the target switch, this is analogous to performing a delayed reach from a suboptimal initial condition. The network will achieve the correct end behavior, because when the go cue is delivered, it must settle to the single stable attractor region of the movement dynamical system, as shown in Fig 7f-i. However, there is the potential for initial aberrant kinematic activity from initiating the movement dynamical system from the fixed point of an incorrect target plan. We found that this was not the case, and indeed the RNN was able to carry out the target switch task with no additional delay period, illustrated by representative examples in Fig 8a. Thus, the mechanism used by the RNN to perform the delayed reach task generalizes to perform two variants of the target switch task, even though it was not trained on it. Analogous arguments apply to the pulsed RNN. We found that, like the sustained RNN, the pulsed RNN also generalized to the target switch task when the switch was delivered either at the same time or preceding the go cue, as shown in Fig 7a.

**Figure 8:**
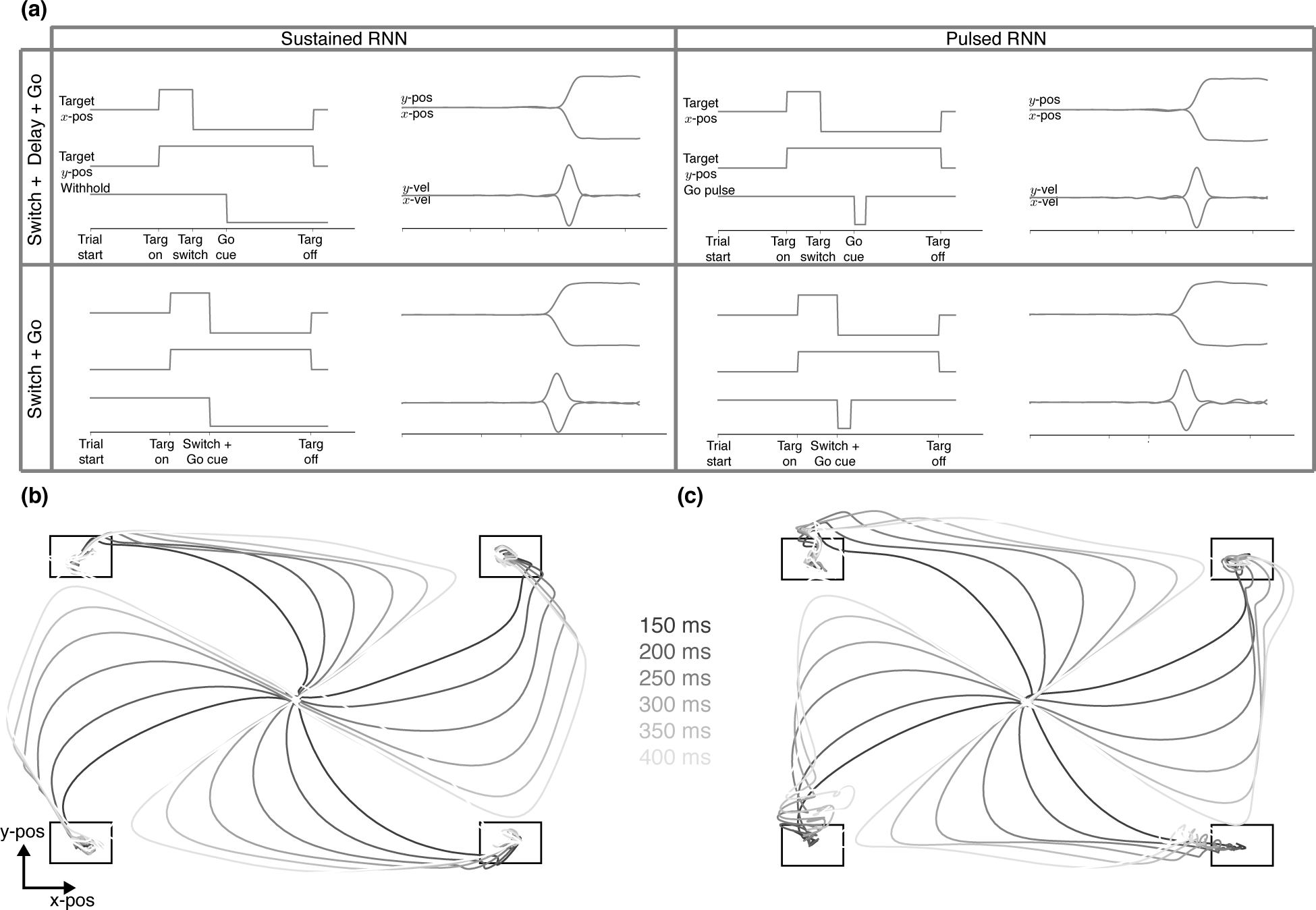
Example behavior for the RNNs generalizing to perform the target switch task. **(a)** Both RNNs discussed generalize to the target switch task when the target switch occurred either before or at the same time at the go cue. Both RNNs had a coefficient of determination *R*^2^ > 0.99 in reconstructing the output kinematics despite not being trained on the target switch task explicitly. **(b)** RNN output positions during a target switch task, where the switch is delivered after the go cue. Here, we show four target switch conditions, when switches occur to adjacent targets. This output corresponds to the RNN trained to perform the sustained go cue delayed reach task. The different shades correspond to output reach trajectories when the target was switched some amount of time following the go cue (legend in between panels (a) and (b)). **(c)** Same as **(b)** but for the pulsed RNN trained on the pulsed go cue delayed reach task.

We also considered a task when the target is switched after the go cue. This task comprises an online corrective component, where feedback of the arm’s kinematics play an important role in updating motor commands to reach to a new target. While we did not account for this corrective component, we nevertheless found that the open-loop RNN could reasonably perform the task as the correct target input dictated the stable attractor location. This is shown in Fig 8b,c, where for different lengths of time after the go cue, a target switch is delivered in the middle of the trial. We found that even though there was not a corrective feedback component and the target was abruptly changed, the RNN made a smooth and reasonable trajectory between targets.

We found that the pulsed RNN had poorer generalization in the presence of input and recurrent noise. We incorporated input noise and recurrent noise into RNNs as they performed a task where the target switched 200 ms after the go cue. We found that, in general, the pulsed RNN had poorer robustness to both input noise and recurrent noise (Fig 9a,b) across varying levels of noise. In particular, we found that the pulsed RNN especially performed worse when the target switch was maximal at 180° (purple lines in Fig 9a,b). The sustained RNN adequately performed this task, being able to make diagonal corrections, as shown in Fig 9c. However, we found that the pulsed RNN was not able to consistently perform this task (example trajectories in Fig 9d). One reason for poorer performance is that on several occasions, the output trajectories began to correct in the right direction, but do not cross the bifurcation axis, settling back to the preparatory state attractor. These results suggest that, while both the sustained and pulsed RNNs are capable of using their dynamical mechanisms to generalize to target switch tasks, the sustained RNN has more robust generalization in the presence of noise.

**Figure 9:**
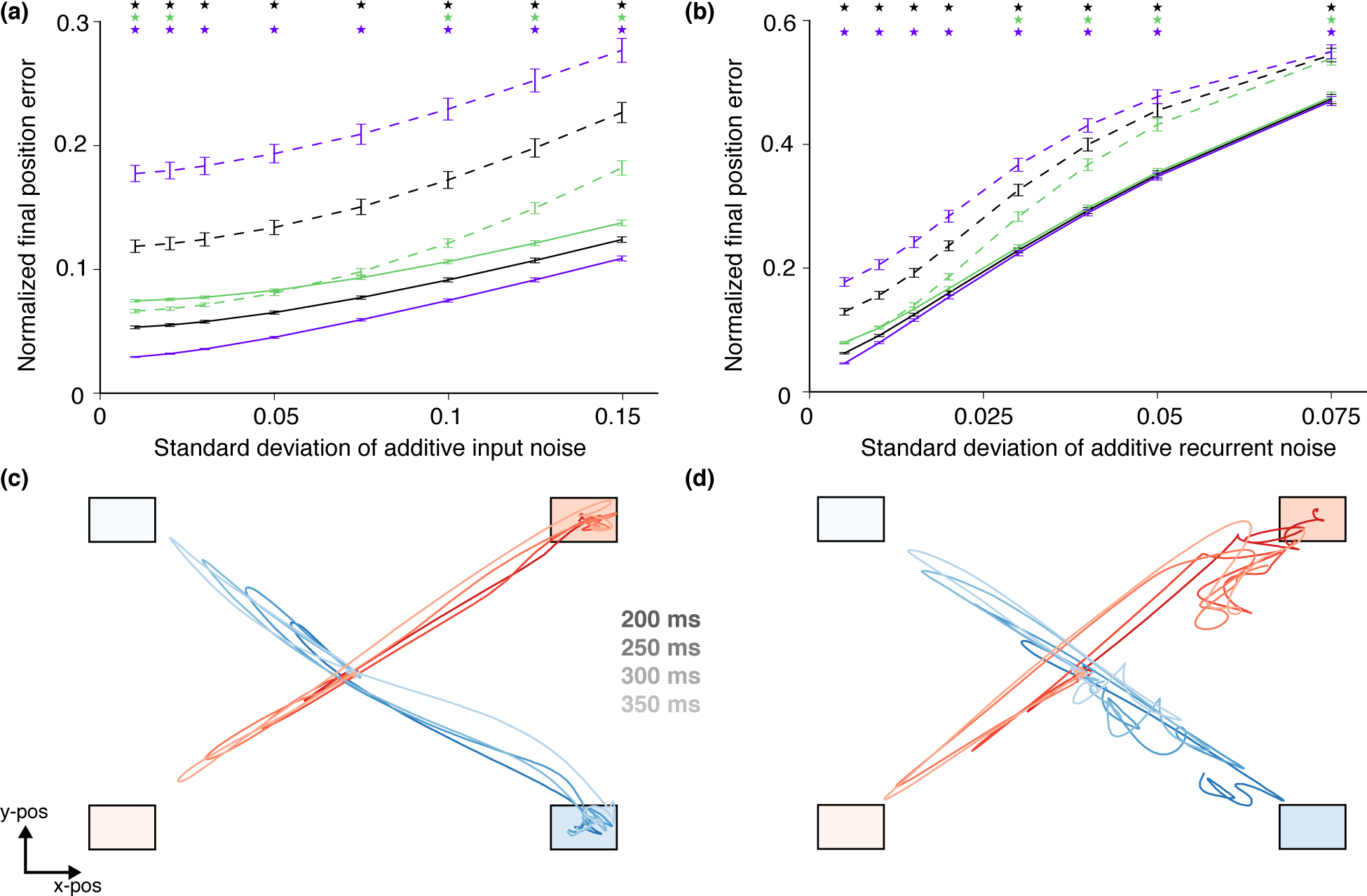
**(a)** Normalized error in the final position as a function of the standard deviation of Gaussian noise added to the input. The final position was measured at 1200 ms after go cue onset, when the trials were terminated. The position errors are normalized so that the target position has a distance of 1. The solid lines correspond to the performance of the sustained RNN trained on the delayed reach task when performing the target switch after go cue task, and the dotted lines correspond to the performance of the pulsed RNN. The black lines denote the error across all switch conditions. The purple lines denote the error for switch conditions where the switched target was 180° away from the pre-switch target, and the green lines correspond to the error for switch conditions where the switched target was +90° away from the pre-switch target. Stars denote a significant difference in the means at the level *p* < 0.01 (bootstrap, 1001 shuffles). Error bars are standard error of the mean. In general, the pulsed RNN has poorer robustness under additive input noise. **(b)** The same as **(a)** but for recurrent noise added to each artificial neuron. In general, the pulsed RNN has poorer robustness under additive recurrent noise. **(c)** Example output kinematics of the sustained RNN for target switch trials for two conditions, where the target switch is diagonal. The lighter target corresponds to the initially prompted target, and the darker target corresponds to the switched target. To make the task harder, noise was injected into the inputs. The RNN arrives at the correct final behavior, as would be expected by its dynamical mechanism. **(d)** Same as **(c)** but for the pulsed RNN trained to perform the pulsed go cue delayed reach task. One can observe that the RNN fails to perform the task adequately, achieving the incorrect final position.

## Discussion

RNNs, being an *in silico* circuit model that is fully observed, are a promising tool to investigate mechanisms underlying motor behavior (Sussillo et al., 2015; Hennequin et al., 2014; Michaels et al., 2016), decision-making (Mante et al., 2013; Chaisangmongkon et al., 2017; Song et al., 2017), and other neurophysiological tasks (Song et al., 2016). However, there are important considerations when using RNNs to make conclusions about cortical computation. Our results highlight two key points: (1) that it is not sufficient to demonstrate that RNNs have artificial neurons that only recapitulate key motifs in single neurons, and (2) that even networks that capture both single neuron and population motifs in the data may use fundamentally distinct mechanisms.

First, we found that it is insufficient to have RNNs merely recapitulate single neuron motifs alone. In addition to single neuron motifs, RNNs ought capture population level motifs in the data. This is clear in the choice of activation function to train RNNs. In RNNs trained with the relu activation, it was possible to find artificial neurons with preparatory activity, but these RNNs did not capture key population features from the data. Rather, preparatory trajectories were relatively short compared to movement trajectories (Fig 4c), which is inconsistent with empirical results. Further, even for the tanh activation function, we found that it was important to regularize the network appropriately to capture preparatory variability in the population. This would not have been straightforward if only looking at single neuron comparisons.

Second, we saw that two distinct RNNs could both recapitulate key hallmarks of neurophysiological activity, but do so in fundamentally different ways. In this manner, even if an RNN recapitulates both single neuron and population level motifs, careful consideration should be given to how the RNN dynamically performs the task. Our results showed that by varying how the task inputs are designed, the RNN can use distinct mechanisms with important consequences on generalization in the presence of noise. Our results also demonstrate that in addition to regularizations (e.g., (Sussillo et al., 2015)), architectures (e.g., (Song et al., 2016)), and training rules (e.g., (Miconi, 2017; Song et al., 2017)), task input design can have an important effect on how the RNN’s computations are performed. It may be possible for future experiments to be designed in such a way to militate between these mechanisms by assessing behavior in the presence of noisy inputs.

As task design can substantially affect the dynamical mechanisms employed by the RNN, so too may other hyperparameters and training paradigms. It will be appropriate to consider how RNN architectures and parameters affect dynamical mechanisms, their robustness to noise, and generalization. For example, Song and colleagues demonstrated that it is possible to design RNNs that obey biological constraints such as Dale’s law (Song et al., 2016). They demonstrated that in these networks, one could find single artificial neurons consistent with empirical recordings. Similarly, Miconi and colleagues demonstrated that networks can be trained with biological learning rules (Miconi, 2017). These learning rules, which reproduce neurophysiological features of the data, may affect the network dynamics. Assessing the extent to which important features of the employed dynamical mechanisms change through introducing biological constraints and learning may play an important role in proposing mechanisms for cortical computation and making concrete predictions for future experiments.

Interrogating an RNN’s dynamics also has consequences for what type of dynamics may be *sufficient* to carry out a class of tasks. In our work, we found that an RNN trained to only perform a delayed reach task was capable of generalizing to target switch tasks, even though it wasn’t trained on these tasks. This shows that the mechanism employed by the RNN to perform a delayed reach task endows the network with the capability of performing the target switch task. An interesting line of future work may assess how the RNN’s dynamics change as it is trained to perform a wider assortment of tasks (Yang et al., 2017). This may describe how many, and what classes, of tasks are necessary to provide an RNN with the capability of performing a different set of tasks. Further, in so far as the capability to perform a variety of tasks changes the dynamical mechanisms of the network, this may help to narrow the set of plausible mechanisms used to perform a given task. For example, if we trained the networks in this work to perform motor tasks with perturbations to the arm (e.g., Omrani et al. (2014); Nashed et al. (2012)), does the network cease to employ a bistable mechanism? Another line of inquiry is to visualize the dynamics of RNNs that fail to generalize, and determine what deficiencies result in poor generalization.

## Acknowledgments

We thank Chandramouli Chandrasekaran and Krishna Shenoy for helpful discussions. To train RNNs, we used pycog (https://github.com/xjwangiab/pycog/) with modifications for task design, architecture and optimization.

## Author contributions

JCK was responsible for design, analyses, simulation experiments, and manuscript writeup.

## Supplementary materials

### Supplementary Movie 1.

Movie demonstrating visualized dynamics at each point in time as the RNN performs a delayed reach task. The stable fixed point jitters in the preparatory period due to numerical precision; we performed the optimization to find the fixed point at each time step. A link to this movie is at: https://seas.ucla.edu/~kao/vid/18rnn_SuppMovie1.mp4

### Supplementary Table 1.

**Table S1:** Parameters used for RNN training. The λ_Ω_ regularizer is described further in the study by (Pascanu et al., 2012) as well as (Song et al., 2016).

| RNN parameters | Values |
| --- | --- |
| Number of units | 100 |
| Time constant ($\tau$) | 50 ms |
| Discretization bin width | 10 ms |
| $L_2$ regularizer for $\mathbf{W}_{\text{in}}$ | $1 \times 10^{-3}$ |
| $L_2$ regularizer for $\mathbf{W}_{\text{out}}$ | $1 \times 10^{-3}$ |
| $L_2$ regularizer for $\mathbf{W}_{\text{rec}}$ | $1 \times 10^{-3}$ |
| $L_2$ regularizer for $\mathbf{r}(t)$ | $1.9 \times 10^{-3}$ |
| $\lambda_{\Omega}$ regularizer | 2 |
| Activation function | $\tanh(\cdot)$ |
| Initial learning rate (Adam) | $1 \times 10^{-4}$ |
| Maximum gradient norm | 0.2 |

### Supplementary Figures.

**Figure S1:**
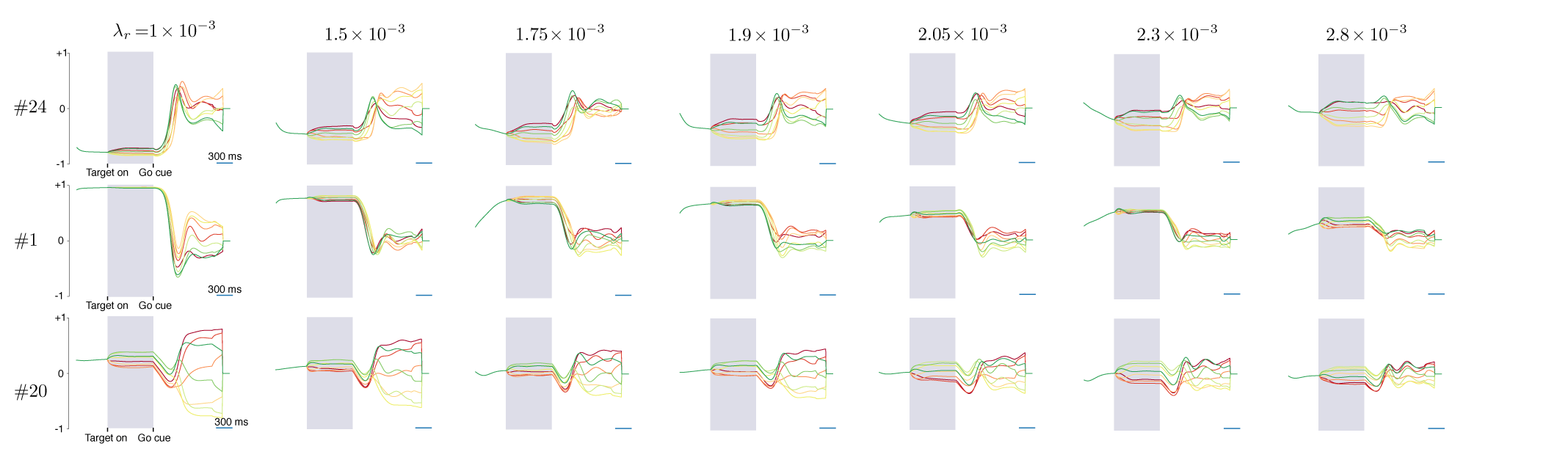
PSTHs for the same neuron in an RNN across seven different levels of regularization for a tanh RNN. The region highlighted in gray corresponds to preparatory activity. The neuron is the “same” across all networks in the sense that we initialized the networks in the exact same way, with the same random seed, and they only differed in the amount of rate regularization. We found that each unit across the different RNNs shared similar motifs under this training process. In general, as rate regularization increases, the units have more preparatory activity relative to movement activity.

**Figure S2:**
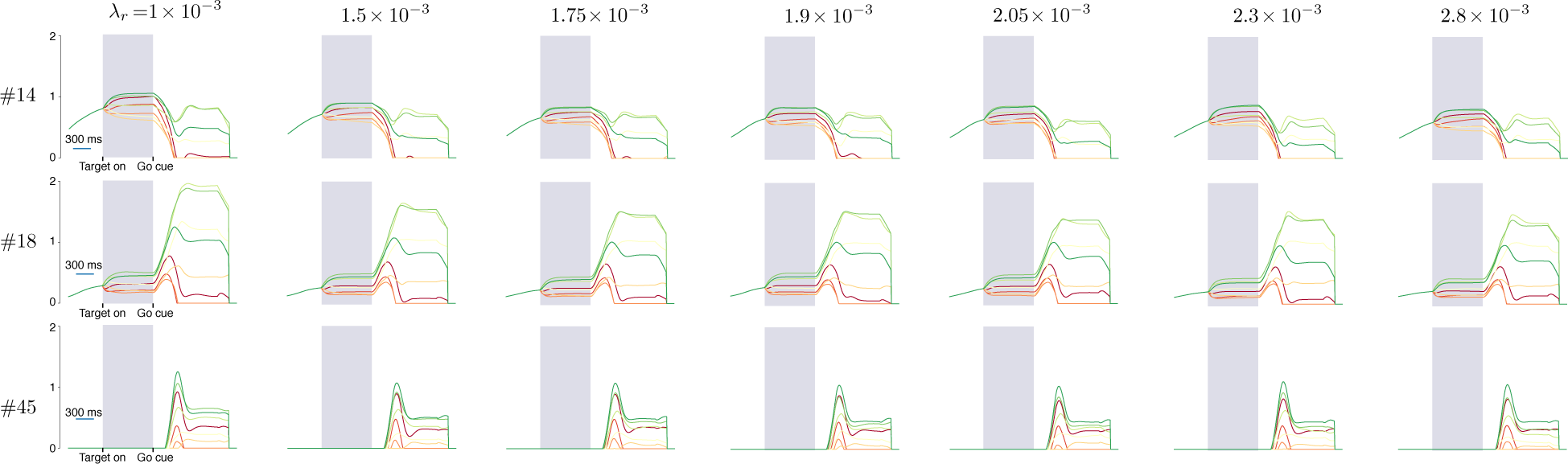
PSTHs for the same neuron in an RNN across 8 different levels of regularization for a relu RNN. The region highlighted in gray corresponds to preparatory activity. In general, even as rate regularization increases, the units have similar activity.

**Figure S3:**
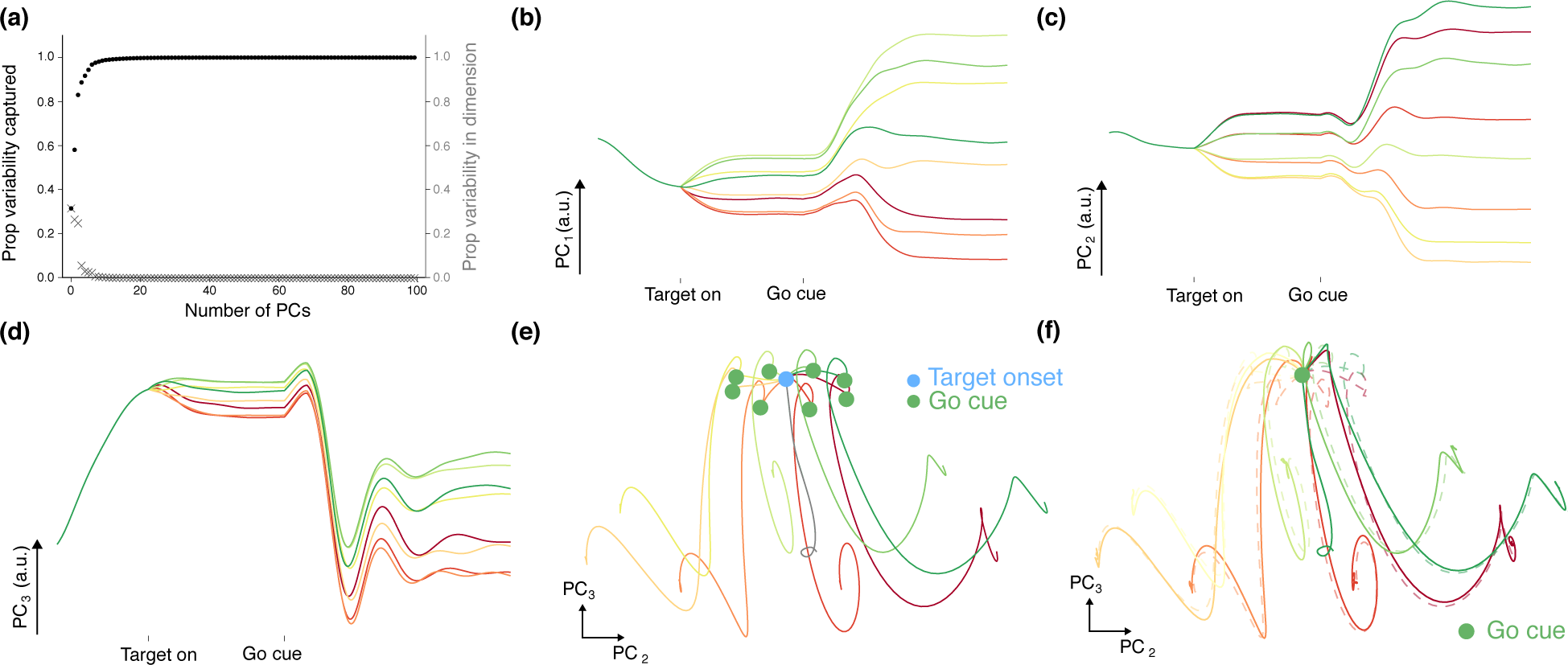
Principal components analysis for RNN trained to perform a delayed reach task with pulsed go cue. **(a)** Variance captured by dimension. The first 5 PCs capture 91.8% of the PSTH variability and the first 10 PCs capture 98.6% of the PSTH variability. **(b)** PC_1_ of the RNN rates. PC_1_ contains a substantial amount of condition dependent information. **(c)** PC_2_ of the RNN rates. **(d)** PC_3_ of the RNN rates. This dimension captures a large transient signal that is largely condition independent after the go cue. **(e)** Projection of PC_2_ and PC_3_ with a delay period. The trajectories in the delay period reach a target dependent stable region in state space, and subsequently are strongly driven along trajectories associated with movement production. **(f)** Projection of PC_2_ and PC_3_ shows that the delay period is not obligatory. The dotted traces are trajectories with a delay period while the solid traces are trajectories without a delay period.

